# Soil depth gradients in microbial growth kinetics under deeply- vs. shallow-rooted plants

**DOI:** 10.1101/2021.04.26.441349

**Authors:** Kyungjin Min, Eric Slessarev, Megan Kan, Karis McFarlane, Erik Oerter, Jennifer Pett-Ridge, Erin Nuccio, Asmeret Asefaw Berhe

## Abstract

Climate-smart land management practices that replace shallow-rooted annual crop systems with deeply-rooted perennial plants can contribute to soil carbon sequestration. However, deep soil carbon accrual may be influenced by active microbial biomass and their capacity to assimilate fresh carbon at depth. Incorporating active microbial biomass, dormancy and growth in microbially-explicit models can improve our ability to predict soil’s capacity to store carbon. But, so far, the microbial parameters that are needed for such modeling are poorly constrained, especially in deep soil layers. Here, we investigated whether a change in crop rooting depth affects microbial growth kinetics in deep soils compared to surface soils. We used a lab incubation experiment and growth kinetics model to estimate how microbial parameters vary along 240 cm of soil depth in profiles under shallow- (soy) and deeply-rooted plants (switchgrass) 11 years after plant cover conversion. We also assessed resource origin and availability (total organic carbon, ^14^C, dissolved organic carbon, specific UV absorbance, total nitrogen, total dissolved nitrogen) along the soil profiles to examine associations between soil chemical and biological parameters. Even though root biomass was higher and rooting depth was deeper under switchgrass than soy, resource availability and microbial growth parameters were generally similar between vegetation types. Instead, depth significantly influenced soil chemical and biological parameters. For example, resource availability, and total and relative active microbial biomass decreased with soil depth. Decreases in the relative active microbial biomass coincided with increased lag time (response time to external carbon inputs) along the soil profiles. Even at a depth of 210-240 cm, microbial communities were activated to grow by added resources within a day. Maximum specific growth rate decreased to a depth of 90 cm and then remained consistent in deeper layers. Our findings show that > 10 years of vegetation and rooting depth changes may not be long enough to alter microbial growth parameters, and suggest that at least a portion of the microbial community in deep soils can grow rapidly in response to added resources. Our study determined microbial growth parameters that can be used in microbially-explicit models to simulate carbon dynamics in deep soil layers.

## Introduction

Soil contains the largest carbon (C) pool in terrestrial ecosystems (IPCC, 2013; Jobbágy and Jackson, 2000). Accrual of organic C along soil depth gradients can contribute to climate change mitigation through sequestration of atmospheric carbon dioxide (Paustian et al., 2016). However, most soil C dynamics research so far has focused on the top 30 cm of soil, where soil C concentrations are highest (Jobbágy and Jackson, 2000; Kögel-Knabner et al., 2008). Recent studies highlight that surface soil (< 30 cm) and deep soil (> 30 cm) C pools respond differently to changes in environmental conditions (Berhe et al., 2008; Fierer et al., 2003a; Jia et al., 2019; Min et al., 2020; Pries et al., 2017). At the surface, variation in abiotic factors such as temperature and moisture can influence microbial transformation of soil organic C. However, in deeper soil layers, relatively constant environmental conditions, increased mineral interactions, and lower microbial biomass, combined with distinct microbial community composition, may alter the capacity for microbial C transformation relative to the surface (Rumpel et al., 2012; Rumpel and Kögel-Knabner, 2011). Given that most soil C is stored in deep soils (Jobbágy and Jackson, 2000), we must establish a better understanding of how microbially-driven C dynamics vary along soil depth gradients.

Microorganisms modify the amount and chemical composition of soil organic matter via decomposition, respiration, growth, and death. Due to the critical roles microbes play in soil C dynamics, several biogeochemical models have recently incorporated microbial parameters (Allison et al., 2010; Salazar-Villegas et al., 2016; Wang et al., 2015; Wieder et al., 2015, 2013). Active microbial biomass and microbial growth rates are both key parameters in microbially-explicit models (He et al., 2015; Zha and Zhuang, 2020), but are poorly quantified because of the technical difficulty of direct quantification (Geyer et al., 2016). Most empirical measurements of microbial biomass reflect total biomass, using analyses made with chloroform fumigation (Anderson and Domsch, 1978; Vance et al., 1987), phospholipid fatty acid (Kao-kniffin and Balser, 2008; Kohl et al., 2015) or ATP (Contin et al., 2001; Martens, 1995) approaches. However, since total microbial biomass comprises both active and dormant biomass, and active microbial biomass is more relevant to biogeochemical cycles (Couradeau et al., 2019; Geyer et al., 2016; Salazar-Villegas et al., 2016; Singer et al., 2017), using total microbial biomass in models may be inadequate when projecting soil C dynamics. Microbial growth and dormancy can be estimated from growth kinetics models (Mitchell et al., 2004; Panikov and Sizova, 1996), as well as quantitative stable isotope probing (Koch et al., 2018; Pett-Ridge and Firestone, 2017; Schwartz et al., 2016), nucleotide analog tagging (Allison et al., 2008; Hanson et al., 2008), and amino acid-tagging approaches (Couradeau et al., 2019; Hatzenpichler et al., 2014).

In 1995, Panikov (1995) developed a model that parses microbial respiration sources into growth vs. non-growth related respiration and estimates growth-related parameters (e.g., active vs. dormant microbial biomass, the lag phase before exponential growth, and maximum specific growth rate; See Methods). This approach is useful because it requires readily measurable microbial respiration as an input, and the estimated microbial parameters can be directly used in microbially-explicit models. Using the Panikov model, Blagodatskaya et al. (2014) demonstrated that rhizosphere microbes exhibit a shorter lag time than non-rhizosphere microbes, due to their higher proportion of active biomass and associated lower dormancy. Also, Salazar-Villegas et al. (2016) demonstrated that neither warming nor wetting treatments alter maximum specific growth rates of soil microbial communities.

Changes in resource availability with soil depth may influence microbial growth parameters. The amounts of organic C, nitrogen (N), and other resources are often higher at the soil surface where inputs from plant litter and root exudates are concentrated (Jobbágy et al., 2001; Rumpel and Kögel-Knabner, 2011). In contrast, deeper layers receive limited direct plant inputs; typically, only dissolved organic matter that moves downward through mass flow or diffusion. In addition, mineral-organic matter associations in deeper layers can reduce microbial access to resources (Schmidt et al., 2011). Some studies suggest that soil microorganisms grow faster when resources are more abundant. For example, Blagodatskaya et al. (2010) demonstrated that more C inputs into soils under elevated CO_2_ increased microbial maximum specific growth rate.

Similarly, rhizosphere microbes exhibited greater maximum specific growth rates and shorter lag times than bulk soil microbes (Blagodatskaya et al., 2014). Recent research suggests that these rapid growth responses to resource additions may be driven by traits related to microbial evolutionary history, taxonomy, 16S rRNA gene copy number or genome size (Li et al., 2019; Morrissey et al., 2019). Given the expected decreases in resource availability along soil depth profiles, it is plausible that microbial dormancy increases and the capacity for a rapid growth response declines with soil depth. It is also possible that microbial lag time increases with soil depth because microbes in deeper layers may be more likely to use strategies such as sporulation, and some of the abundant taxa in deep soils have been shown to decrease in relative abundance after soil fertilization (Brewer et al., 2019). Yet, it is unclear how the direction and magnitude of microbial growth parameters vary with soil depth profiles.

Improved understanding of depth-specific microbial growth kinetics is key for climate-smart land management practices that consider implementation of improved root phenotypes and/or replacement of annual crops with deeply rooted perennials (Paustian et al., 2016). One model deeply rooted species that is being considered in these efforts is switchgrass, a perennial grass native to North America, with a deep rooting system that extends up to 3 m (Liebig et al., 2005; Wright, 2007). Conversion from annual crops to switchgrass can significantly increase deep soil C stocks (Slessarev et al., 2020b). Deeply rooted plant contributions to increased soil C storage are likely tied to changes in the distribution of rhizodeposits, water flow through the soil profile, and the types of microbial communities that thrive in the rhizosphere. For example, in switchgrass fields, enhanced production of extracellular polymeric substances and higher soil aggregate formation have been measured relative to annual crops (Sher et al., 2020), and linked to increased soil C storage with depth (Blanco-Canqui et al., 2005; McGowan et al., 2019; Sher et al., 2020). Also, switchgrass roots increased microbial OTU (operational taxonomic unit) richness along 60 cm of the soil profile (He et al., 2017). To our knowledge, no data currently exists on how the conversion of an annual crop to a deep-rooted perennial crop, such as switchgrass, impacts microbial growth and growth-related parameters (e.g., dormancy, active biomass) in deeper soil layers.

Here, we investigate microbial growth parameters along soil depth profiles under deeply- (switchgrass) and shallow-(soy) rooted plants. We collected soils under switchgrass and soy plots (n=3, each) from adjacent, paired field locations at soil sampling depths ranging from 0 to 240 cm. We incubated these soils with yeast extract as an added C and energy source, and monitored CO_2_ efflux to parameterize a growth kinetics model according to a previously described approach (Blagodatsky et al., 2006; Wutzler et al., 2012). By fitting a growth kinetics model to the CO_2_ data, we estimated total and active microbial biomass, maximum specific growth rate, and lag time. In addition, we quantified resource availability prior to the yeast extract addition to explore the interaction of depth and resource availability on microbial parameters. We hypothesized that (1) increasing depth would decrease maximum specific growth rate, due to decreasing resource availability (available C, N), (2) active biomass would be highest in surface soil layers while lag time would be greatest in deep soil layers, due to reductions in resource availability, and (3) microbial growth parameters would be less affected by depth under switchgrass than soy, because greater rooting depth results in more abundant resources inputs at depth.

## Methods

### Study site and soil sampling

The study site is located near Bristol, South Dakota (45°16’N, 97°50’W), where switchgrass (*Panicum virgatum* L.; cultivar, Sunburst) has been growing since 2008 (Sekaran et al., 2019). Before switchgrass was cultivated, the site was used for soy. The mean annual temperature is 6.3°C, mean annual rainfall is 638 mm, and mean annual snowfall is 1092 mm at the nearest weather station in Webster, South Dakota (∼32 km away, https://www.usclimatedata.com/climate/webster/south-dakota/united-states/ussd0368/2018/1). The study site is part of a long-term nitrogen (N) addition experiment (0, 56, and 112 kg urea-N ha^−1^ yr^−1^ since 2008), however, only control sites (0 kg N) were used for this experiment. The soils are classified as Fine-loamy, mixed, superactive, frigid Calcic Hapludolls (Barnes series, USDA Soil Taxonomy) and Fine-loamy, mixed, superactive, frigid Typic Calciudolls (Buse series, USDA Soil Taxonomy). The parent material is glacial till dominated by fine-grained sedimentary rocks deposited in the upper Pleistocene (Mankato substage, approximately 14 ka before present (Flint, 1955)). Each treatment plot is 21.3 m x 365.8 m with a 2-20% slope.

In July 2019, we collected a 240 cm deep soil core under switchgrass from each of three non-fertilized plots on a crest landscape position (n=3) using a Geoprobe® 54LT direct push machine (Salina, Kansas, USA) combined with a MC7 system (diameter: 7.62 cm). For comparison, we also collected three 240 cm of deep soil cores in a soy field adjacent to the switchgrass plots (n=3). Since 2008, this field has been primarily used to grow soy and intermittently cropped with other annuals (spring wheat or corn or maintained as pasture (USDA NASS 2020; https://nassgeodata.gmu.edu/CropScape/)). After extraction, soil cores were immediately divided into nine sections using a hybrid fixed depth and soil horizon method, i.e., when there was a clear separation between soil horizons by color or texture, we divided sections using a generic horizon approach. Otherwise, we divided soil cores into 20 cm increments for the top 60 cm soil and 30 cm increments for depths below 60 cm. Hereafter, we refer to the soil depth sections as 0-20, 20-40, 40-60, 60-90, 90-120, 120-150, 150-180, 180-210, and 210-240 cm for simplicity of visual representation. After soil cores were subsectioned, we removed rocks and roots manually in the field, shipped the samples to the University of California-Merced in a cooler, and stored them at 4°C. Within a week of arrival, soil samples were processed to determine soil physical, chemical, and microbial properties. In the lab, more roots were hand-picked if necessary. Roots collected from soil cores were dried and weighed to quantify root distribution along soil depth profiles.

### Soil physicochemical properties

Soil pH was determined in water (1:5, fresh soil weight:water volume) using a Mettler Toledo pH meter. We extracted dissolved organic C (DOC) by mixing 6 g of fresh soil with 30 mL of 0.5 M K_2_SO_4_ and shaking the soil solution for 4 h. The soil solution was then centrifugated at 2,000 rpm for 5 min and the supernatant was filtered through a 0.45 µm membrane filter (PALL, Port Washington, NY, USA). The filtrate was stored at −20°C until analysis. The DOC concentration and dissolved N in K_2_SO_4_ extracts were determined using a VCSH Total OC Analyzer (SHIMADZU, Kyoto, Japan) after thawing in the Environmental Analytical Lab at the University of California-Merced. To estimate the easiness of microbial utilization, we quantified Specific UV Absorbance at 254 nm (SUVA) on K_2_SO_4_ extracts, using a UV-VIS spectrophotometer (Evolution 300, Thermo Scientific, Massachusetts, USA). Higher SUVA values indicate a higher relative abundance of aromatic compounds in DOC extracts (Weishaar et al., 2003) and less available form of DOC to microbes.

Total C and N were quantified on dried and ground soils at the Oregon State University Crop and Soil Science Central Analytical laboratory. Because soils in both vegetation types were alkaline with pH between 8.0 and 9.3 (S1), we treated soils with 1M HCl to remove carbonates before determining total OC concentrations. Inorganic C was quantified at Lawrence Livermore National Laboratory by treating finely-ground subsamples of each sample with 1 M phosphoric acid in a sealed jar and measuring CO_2_ evolved using a LI-850 infrared gas analyzer (Robertson et al., 1999) for 24 h. Total OC and ^13^C were quantified on acid-treated soils (treatment with 1 M HCl) at the Integrative Biology Center for Stable Isotope Biogeochemistry Lab at the University of California-Berkeley.

We assessed radiocarbon values on soils from 0 cm to 150 cm only. Radiocarbon values were measured on the NEC 1.0 MV Tandem Accelerator Mass Spectrometer (AMS) or the FN Tandem Van de Graaff AMS at the Center for Accelerator Mass Spectrometry (CAMS) at Lawrence Livermore National Laboratory. Prior to measurement, acid-treated soils were prepared for ^14^C measurement by sealed-tube combustion to CO_2_ in the presence of Ag and CuO and reduced onto Fe powder in the presence of H_2_ (Vogel et al., 1984). The ^14^C content of each sample was reported in Δ^14^C notation, corrected for mass-dependent fractionation with measured δ^13^C values, and then corrected to the year of measurement for ^14^C decay since 1950 (Stuiver and Polach, 1977).

### Soil incubation and CO_2_ measurement

For soil incubations, 20 g of fresh soil was weighed into a polypropylene incubation jar (McMaster-Carr, 473 mL) and pre-incubated at 22°C overnight (108 jars = 2 vegetation types * 9 depths * 3 replicate cores * 2 amendment treatments). For treatment jars, we added 40 mg of powder yeast extract g^−1^ dry soil as a C and nutrient source (VWR). We chose yeast extract over glucose to provide more varied forms of C and thus stimulate more diverse microorganisms (Fierer et al., 2003b; Slessarev et al., 2020a). A preliminary experiment confirmed that the amount of yeast extract added to the soil was enough to avoid limiting growth during ∼10 h of exponential growth phase. After the yeast extract was added, we adjusted the soil water content to 60% of water holding capacity (initial gravimetric water content was 18.6 ± 3.7%) and stirred the soils for 30 s. For control jars, only water was added to soils without yeast extract. The jars were closed with a lid equipped with a non-dispersive infrared CO_2_ sensor (CM0126-FS, 1% CO_2_ sensor, CO_2_ meter) (Harmon et al., 2015). We monitored CO_2_ concentrations for 10 min every 30 to 60 min during the incubation (24∼30 h), computed the slope of CO_2_ concentration over time, and quantified the microbial respiration rate (µg C-CO_2_ g^−1^ dry soil h^−1^).

### Microbial growth kinetics

We measured microbial growth kinetics using the Panikov model approach, which makes three assumptions (Panikov, 1995; Panikov and Sizova, 1996): First, microbes are not limited by resources (C, water, and nutrients) during incubation. Second, microbial parameters that are estimated from the model refer to those for the initial microbial community, not for the final microbial community after the activation by resources. Three, microbial parameters estimated from the model refer to those from a whole microbial community, undifferentiated among microbial taxa. Under such conditions, microbial respiration follows the equation:

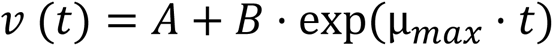

Where *v* is total microbial respiration rate (µg C-CO_2_ g^−1^ dry soil h^−1^) at time *t*, *A* is non-growth respiration of the initial microbial community (µg C-CO_2_ g^−1^ dry soil h^−1^), *B* is growth-based respiration of the initial microbial community (µg C-CO_2_ g^−1^ dry soil h^−1^), *µ_max_* is the maximum specific growth rate of the initial microbial community (h^−1^), and *t* is the time elapsed since the C and nutrients addition (h). Fitting of this growth kinetics model to our CO_2_ data was restricted to the initial exponential growth phase (inflection point) to accurately capture unlimited growth and maximize the goodness of fit *r^2^* (Wutzler et al., 2012).

Physiological status of the initial microbial biomass (relative active biomass) was calculated as below.

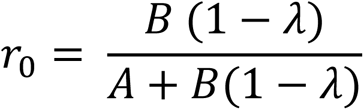

Where *r_o_* is relative active biomass of the initial microbial community (unitless) and *λ* is the ratio of maximum specific rate of growth-related substrate uptake over maximum specific rate of total substrate uptake under unlimited growth. Trutko et al. (1984) demonstrated that the value of *λ* varied between 0.8-0.9 over 100 microbial species. As previous studies exploring microbial growth kinetics have employed *λ* =0.9 (Blagodatskaya et al., 2014; Blagodatsky et al., 2006; Panikov and Sizova, 1996), we also used 0.9 for this study. The value of *r_o_* varies between 0 and 1, with 0 when all microbes are dormant and 1 when all microbes actively grow and divide cells.

Total initial microbial biomass was calculated as per the following:

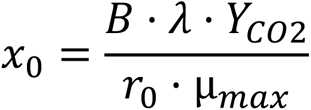

Where *x_o_* is the total initial microbial biomass (µg C-biomass g^−1^ dry soil), and Y_CO2_ is biomass yield per unit CO_2_ respired. We acknowledge that Y_CO2_ can vary with changes in environmental conditions such as temperature and C availability (Lehmeier et al., 2016; Manzoni et al., 2012; Min et al., 2016). However, during the experiment, microbes were allowed to grow under unlimited C and at a constant temperature of 22°C. Thus, we assumed that Y_CO2_ is constant at 1.5, which corresponds to a commonly observed microbial C use efficiency of 0.6 (Keiblinger et al., 2010; Manzoni et al., 2012; Min et al., 2016).

We define lag phase (*t_lag_*) as the period when non-growth respiration is equal to or less than growth-related respiration.

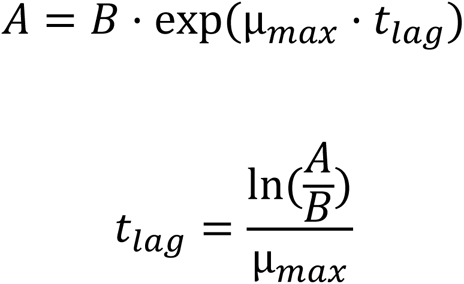

We assume the longer the lag time is, the higher degree of microbial dormancy. Details about the derivatization and calculation of these growth kinetics equations are provided in previous studies (Blagodatsky et al., 2006; Panikov, 1995; Panikov and Sizova, 1996).

### Statistics

Data are presented as mean ± standard error, where n = 3 from replicate cores across soil depth profiles and vegetation types. We fit the Panikov model to the microbial respiration data and estimated microbial growth parameters using a nonlinear least-squares approach (*nls* function in R, R Core Team 3.6.3). The effects of vegetation and depth on microbial growth parameters, resource availability, and root biomass were tested using a linear mixed-effects model, with vegetation and depth as explanatory variables and soil core as a random variable (nlme package, *lme* (linear mixed effects) function, restricted maximum likelihood method). Depth was treated as continuous. When there was a significant interaction of vegetation type and depth, we used Tukey *posthoc* tests to compare soy and switchgrass values at each depth (multcomp package, *glht* (general linear hypotheses) function, Tukey method). All significance was tested at α= 0.05.

## Results

### Depth profiles of root biomass and resource availability

Increasing depth decreased root biomass for both vegetation types (*p* = 0.002; Fig. 1). No root was detected below 60 cm under soy, while we found roots under switchgrass till 150-180 cm.

**Fig. 1.**
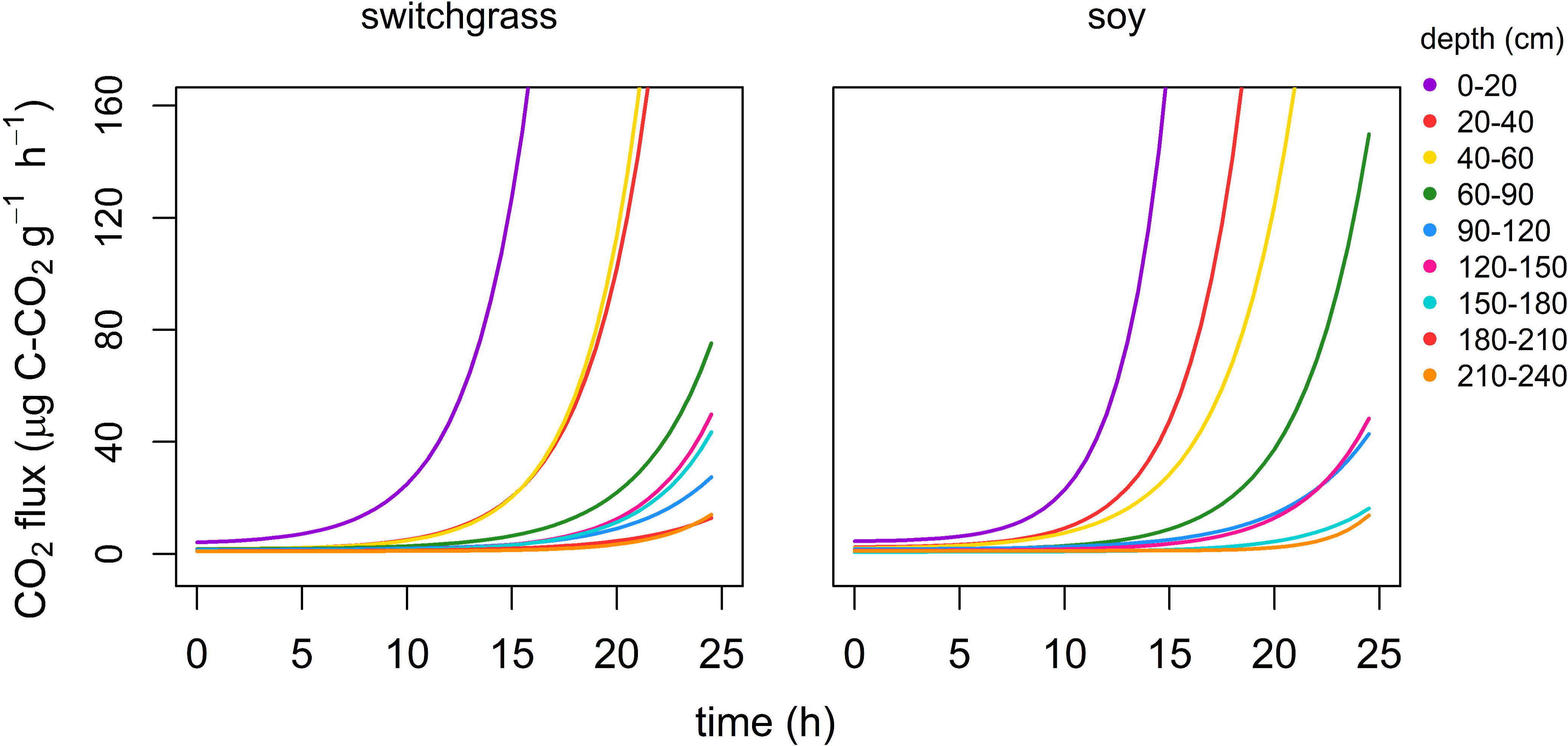
Depth distribution of root biomass under switchgrass (blue) and soy (orange) in clay-loam agricultural fields in South Dakota.

At a given depth interval, switchgrass produced more roots than soy along soil depth profiles (*p* = 0.004; Fig.1).

Overall, resource availability declined with soil depth but did not differ between vegetation types (Fig. 2). For example, total OC was highest at 0-20 cm for both vegetation types, decreased between 0-20 cm and 40-60 cm, then kept similar below (Fig. 2a). We observed greater total OC concentrations under soy than switchgrass at 0-20 cm. Soil depth decreased ^14^C values, but vegetation types had little influence on the ^14^C values (Fig. 2b). Below 90 cm, ^14^C values approached −1000 per mille, which suggests a mean OC age greater than the geologic substrate’s age (Upper Pleistocene ∼ 14 ka (Martin et al., 2004)). This likely reflects contributions of fossil OC from the Pierre Shale, a major constituent of glacial till in eastern South Dakota (Flint, 1955), and the underlying bedrock in the region (Tomhave and Schulz, 2004) with an OC content up to several percent (Gautier, 1986; Kennedy et al., 2002).

**Fig. 2.**
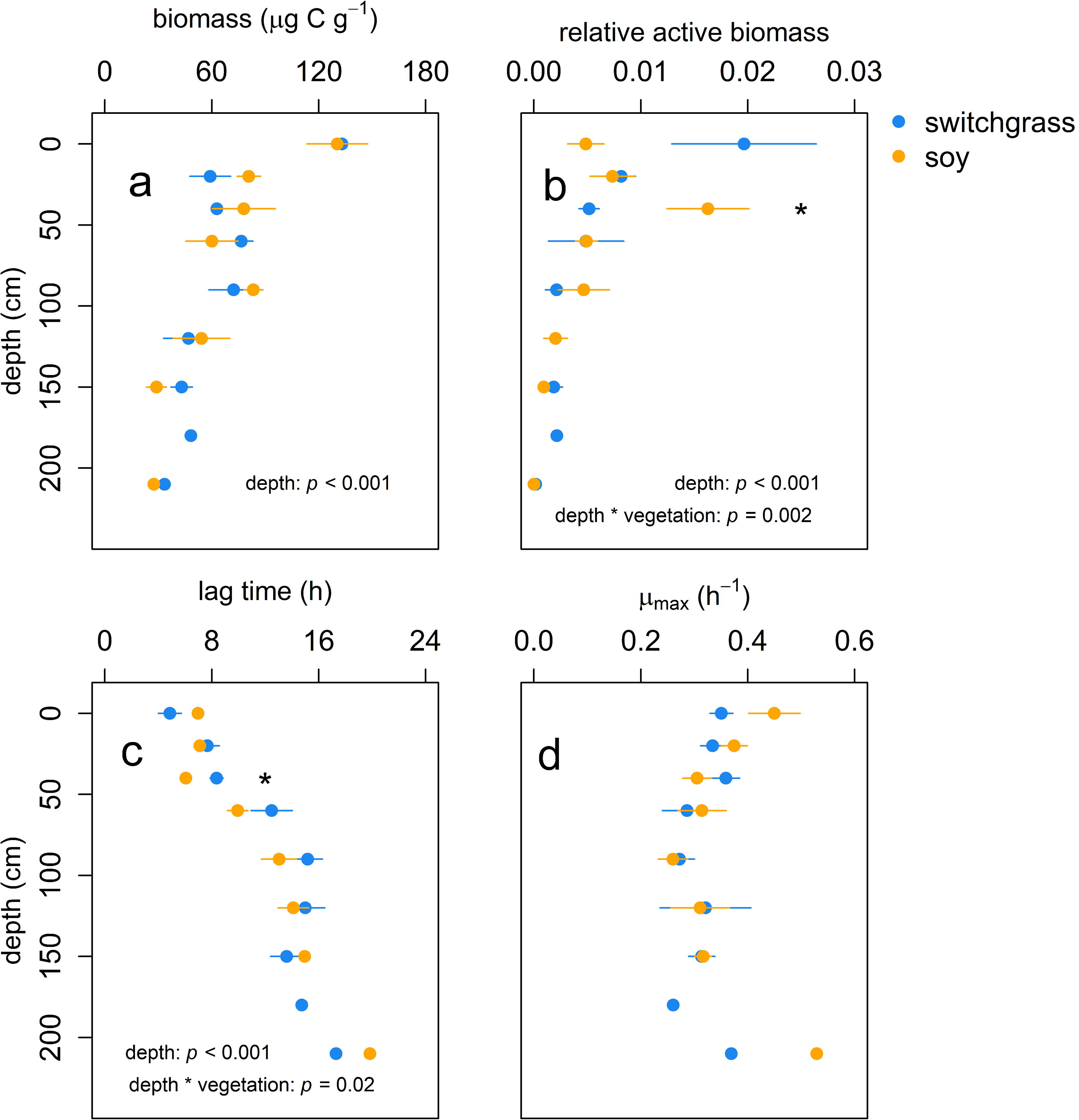
Depth distribution of soil chemistry variables measured in switchgrass (blue) and soy (orange) clay-loam agricultural fields in South Dakota: (a) total organic carbon; (b) ^14^C; (c) dissolved organic carbon; (d) specific UV absorbance of dissolved organic carbon (an index of aromaticity of dissolved organic carbon); (e) total nitrogen; (f) dissolved nitrogen. (n=3, error bars represent ± 1 standard error).

DOC significantly decreased with depth and did not vary between vegetation types (Fig. 2c). At 0-20 cm, the average DOC concentration across vegetation types was 113.2 µg g^−1^, twice that at 20-40 cm (*p* < 0.001). Below 60 cm, the DOC concentration was relatively similar across depths. Depth, but not vegetation type, significantly influenced aromaticity of DOC, measured as SUVA (Fig. 2d).

Both total N and dissolved N were significantly influenced by the interaction of depth and vegetation (*p* < 0.001 and 0.012, respectively; Fig. 2e,f). At 20-40 cm and 40-60 cm, total N content was higher under soy than under switchgrass (*p* = 0.03 and 0.07, respectively). In contrast, dissolved N was similar between soy and switchgrass at 0-20 cm, but was significantly higher under soy at depths below 20 cm.

### Microbial respiration

Basal respiration, before adding yeast extract, was similar across vegetation and depth (S2). When the data was pooled, the average rate of basal respiration was 0.45 ± 0.02 µg C-CO_2_ g^−1^ dry soil h^−1^. We observed distinct patterns in microbial respiration after the addition of water (control) versus water plus yeast extract (treatment) to soils (Fig. 3). Water addition did not alter microbial respiration (Fig. 3a), whereas water plus yeast extract immediately enhanced microbial respiration more than ten times (Fig. 3b). Then, in the following 30 hours, the yeast extract treatment’s respiration rate became relatively stable, increased exponentially, then dropped when all the available resources were consumed.

**Fig. 3.**
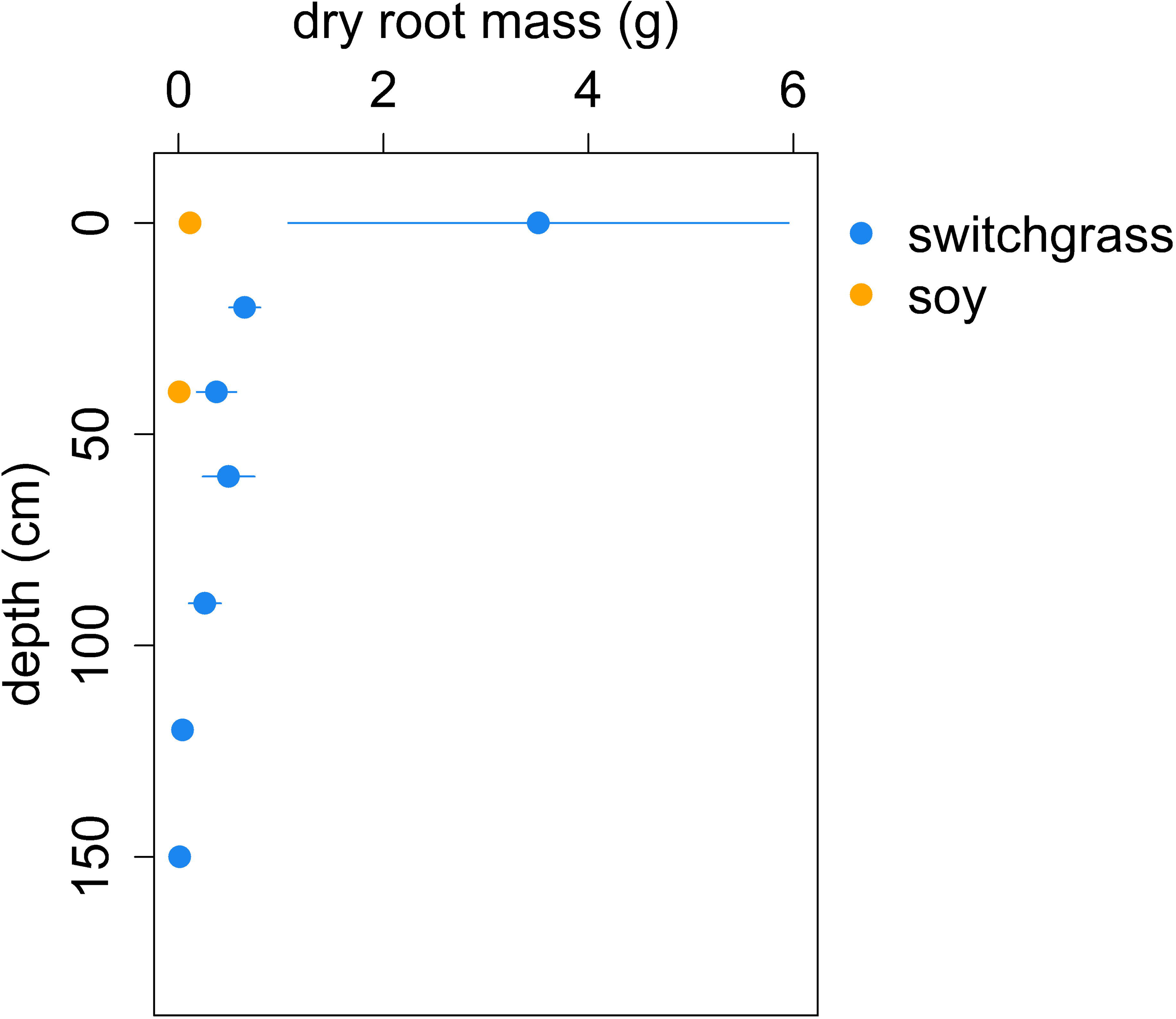
An example plot of soil microbial respiration from an agricultural soil incubated with (a) water only or (b) yeast extract plus water. The red dashed line indicates the time when water or yeast extract plus water was added.

### Growth parameters

We fit the Panikov model to the microbial respiration data to compare microbial growth parameters across depth and vegetation (Fig. 4). Similar to the patterns in resource availability across vegetation and soil depth, we observed a significant depth effect on microbial parameters (except for maximum specific growth rate), but vegetation had little effect on microbial growth kinetics (Fig. 5).

**Fig. 4.**
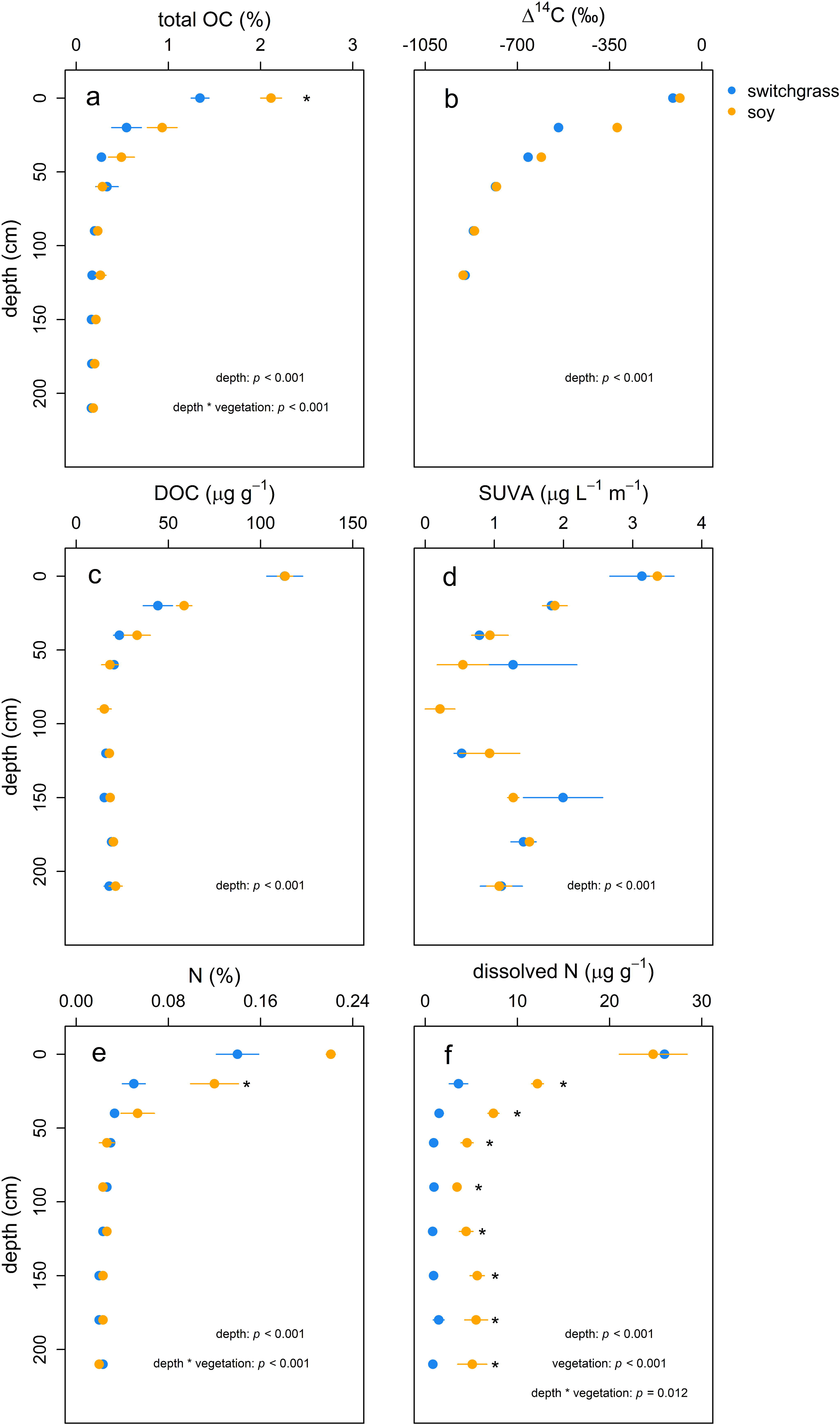
Fitted model of microbial respiration from agricultural soil incubated with yeast extract for switchgrass (left) and soy (right) soils from 0 - 240 cm soil profiles. We used a growth kinetics model described in Panikov (1995). Different colors indicate different depth intervals. For visual simplicity, error bars have been omitted.

**Fig. 5.**
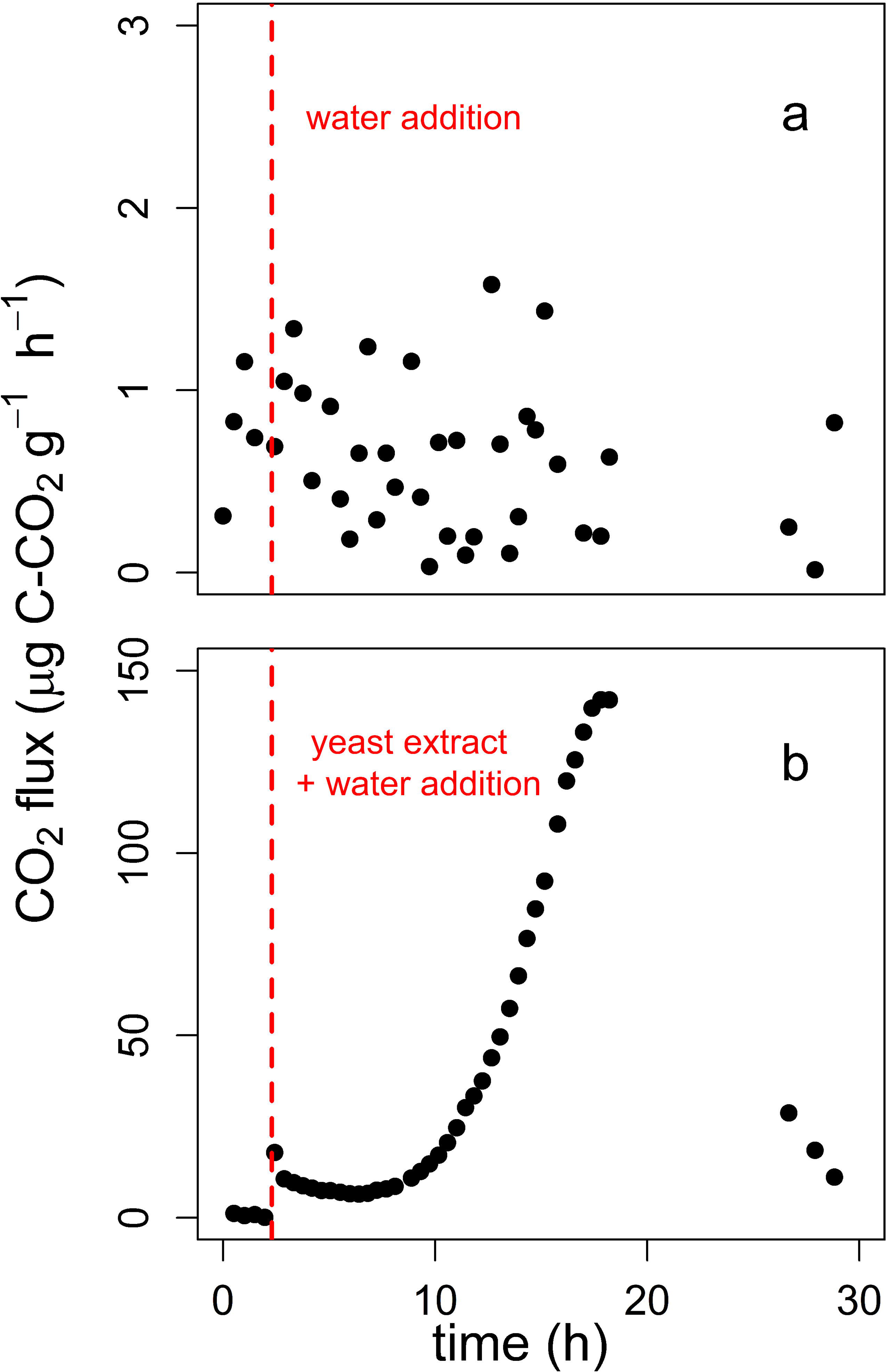
Depth distribution of microbial growth parameters estimated from a growth kinetics model in Panikov (1995), measured in switchgrass (blue) and soy (orange) soils incubated with yeast extract: (a) total microbial biomass before exponential growth; (b) relative active biomass before exponential growth, which varies between 0 and 1. If 0, all the biomass is dormant. If 1, all microbes actively grow and divide; (c) lag time, the response time of microbial community respiration rates to the additions of yeast extract; (d) maximum specific growth rate (µ_max_), the maximum microbial growth potential per unit biomass per unit time (n=3, error bars represent ± 1 standard error).

Total microbial biomass was significantly influenced by depth, but not by vegetation or vegetation and depth interaction (Fig. 5a). When we pooled the data across vegetation, total microbial biomass was 131.8 ± 9.6 µg C g^−1^ dry soil at 0-20 cm and decreased by 77% to 30.5 ± 3.0 µg C g^−1^ dry soil at 210-240 cm.

Overall, the relative active biomass decreased along soil depth profiles, with a 143 times of reduction from the surface to a depth of 210-240 cm (Fig. 5b). We found that at 40-60 cm, where there was no difference in the total microbial biomass between soy and switchgrass (Fig. 5a), the relative active biomass was two times greater under soy than under switchgrass (*p* = 0.04; Fig. 5b).

Lag time increased with soil depth profiles (Fig. 5c) and decreased with relative active biomass (S3). The interaction of depth and vegetation significantly influenced lag time (Fig. 5c). At 40-60 cm, the average lag time under soy was 6.1 h, 27% faster than microbes under switchgrass (*p* = 0.024). At 0-20 cm, the lag time was marginally shorter for microbes under switchgrass than those under soy (*p* = 0.07). We observed that microbes at 210-240 cm were activated to grow within 20 h after the yeast extract additions (Fig. 5c).

Even though the maximum specific growth rate (µ_max_) declined between 0 cm and 90 cm (*p* = 0.011), when we included all depths between 0-240 cm, the depth effect was not significant (*p* =0.234). The average value of maximum specific growth rate was 0.33 ± 0.01 h^−1^ across depth and vegetation (Fig. 5d). The maximum specific growth rate was independent of microbial biomass, regardless of whether we compared it to total microbial biomass (Pearson’s correlation coefficient *r* = −0.41, R^2^< 0.001, *p*=0.01) or relative active biomass (Pearson’s correlation coefficient *r* = −0.18, R^2^= 0.004, *p*=0.28).

Except for maximum specific growth rate, microbial growth parameters had a strong relationship with total OC, DOC, total N, and dissolved N (Table 1). Resource availability was positively related to total and relative active biomass, and negatively related to lag time. The µ_max_ values had no relationship with any soil chemistry index tested in this study.

**Table 1.** Pearson’s correlation coefficient between growth kinetics parameters and soil chemistry.

|  | OC | DOC | SUVA | N | Dissolved N |
| --- | --- | --- | --- | --- | --- |
| Total biomass | **0.47** | **0.46** | 0.09 | **0.49** | **0.41** |
| Relative active biomass | **0.63** | **0.75** | 0.28 | **0.60** | **0.76** |
| Lag time | **-0.70** | **-0.70** | -0.25 | **-0.63** | **-0.59** |
| $\mu_{\max}$ | 0.31 | 0.22 | 0.12 | 0.26 | 0.15 |

Bold when $p < 0.05$

## Discussion

Active microbial biomass drives biogeochemical processes (Couradeau et al., 2019; Geyer et al., 2016; Graham et al., 2016; Salazar-Villegas et al., 2016; Singer et al., 2017; Wutzler et al., 2012), but is hard to characterize due to the inefficiency of methods available to extract biomass, especially from deep soil layers, and the possibility of altering microbial physiological status during extraction (Blagodatskaya and Kuzyakov, 2013a). We reduced this limitation by using a growth kinetics model and examined how depth and vegetation type influence microbial growth potentials. To our knowledge, this is the first study to assess microbial growth parameters along deep soil profiles. Overall, we did not observe significant differences in resource availability between vegetation types (see discussion 3 below), thus we primarily focus on the effects of soil depth on microbial parameters. We demonstrate that microbial communities in deep soil layers can grow relatively quickly in spite of significant evidence of dormancy (lower relative active microbial biomass), and that dormancy becomes more common as soil depth increases.

### 1. Maximum specific growth rate remains unchanged across 240 cm of soil profiles

In this study, maximum specific growth rate was relatively similar across the whole soil profile (0-240 cm; Fig. 5d), yielding an average of 0.33 ± 0.01 h^−1^. Our estimation is comparable to those observed in other studies; maximum specific growth rate was 0.24-0.26 h^−1^ in agricultural topsoils (Blagodatskaya et al., 2014), 0.28-0.29 h^−1^ in soil suspension amended with acetate (Van De Werf and Verstraete, 1987), and 0.25-0.38 h^−1^ in an Ap horizon of loamy soil (Blagodatskaya et al., 2009). We acknowledge that the third assumption of the Panikov model, the whole microbial community behaves as a single entity (see Methods), is likely to be violated in the real world. This is because distinct microbial taxa often exhibit different physiological properties, including growth rate (Ho et al., 2018; Keiblinger et al., 2010; Koch et al., 2018; Morrissey et al., 2019, 2016). As such, we likely overestimated the whole community growth rate, because relatively slowly growing microbes may have not responded to the yeast extract additions and thus we may have only captured the growth of fast growers.

Contrary to our first hypothesis that decreasing resource availability would reduce maximum specific growth rate across the soil profiles, there was no clear relationship between resource availability (C and N) and maximum specific growth rate (Table 1). High resource availability, especially for C, often triggers microbial transition from dormant to potentially active states (Blagodatskaya and Kuzyakov, 2013b; Kovárová-Kovar and Egli, 1998; Lennon and Jones, 2011). Yet, once microbes start growing in response to new resources, antecedent resource availability (i.e., resource availability before adding substrate) may be less relevant to achieving maximum growth rates. Instead, our observation suggests that intrinsic limits of microbial physiology may determine maximum growth potential (e.g., how fast intracellular enzymes catalyze biomolecule synthesis, how constitutively vs. inducibly enzymes are produced)(Adadi et al., 2012), or by the evolutionary history of individual taxa (Morrissey et al., 2019). Although they did not test for maximum specific growth rate, Stone et al. (2014) demonstrated that some extracellular enzymes’ biomass-normalized activity does not change with soil depth despite decreasing resource availability along soil depth gradients. Invariant maximum specific growth rates across depth indicate that microbial communities maintain their growth potential regardless of the environmental conditions they inhabit.

Relatively invariant maximum specific growth rates across the whole soil profile have two ecological implications. First, the potential for microbial C transformation may be comparable between surface and deep soil layers. Most previous studies exploring microbial activity have focused on surface soils, assuming microbial activity is negligible in deeper soil layers. However, our data suggests microbial communities in deep soils can grow and transform soil C as much as surface microbes do. We acknowledge that we have only limited amount of data at 240 cm, but our data suggest that microbial communities at 240 cm exhibited similar maximum specific growth rate to those at surface soils (Fig. 5d). In addition, deep soil microbial communities at 210-240 cm were activated by C addition within 20 h (Fig. 5c) and increased their respiration rate (Fig. 4). Our results are in line with recent findings that microbial C transformations and associated CO_2_ fluxes in deep soils (70 cm) are substantial (Min et al., 2020), microbial activities below 30 cm can be as high (Jones et al., 2018) or higher (Stone et al., 2014) than those at the surface. Also, a recent study revealed that root exudates from switchgrass enhanced microbial production of extracellular polymeric substances and soil C stability along 120 cm of soil profiles (Sher et al., 2020). Thus, given the similar maximum specific growth rates across depth profiles, potential microbial C transformation in deep soil layers can be relatively high.

Second, model representation of microbial growth across soil depth profiles can be simplified. A growing number of studies demonstrate that including microbial parameters improves model projections of the global C budget (Allison et al., 2010; Salazar-Villegas et al., 2016; Wang et al., 2015; Wieder et al., 2015, 2013). However, microbially-explicit models suffer from a lack of empirical knowledge on microbial parameters. Models require information on microbial biomass and biomass-specific rates, as process rates are often expressed as a product of the two (rates = biomass * biomass-specific rates). Our study provides evidence that models may treat microbial growth rate as a constant with soil depth, putting more emphasis on the accurate estimation of microbial biomass. Importantly, it will be critical to quantify active biomass because the active pool of microbes is a better predictor for biogeochemical processes than total biomass (Barnard et al., 2015; Couradeau et al., 2019; Salazar-Villegas et al., 2016; Salazar et al., 2019). In this study, we observed that total biomass was less sensitive to changes in soil depth than relative active biomass. For instance, total biomass decreased by four times while active biomass decreased by 134 times when soil depth changed from 0-20 cm to 210-240 cm (Fig. 5a,b). This suggests that if models employ total biomass, the errors associated with microbial process rates estimation would amplify with soil depth. Empiricists need to use approaches that allow them to distinguish active from total biomass and better inform models of how active biomass would vary when environmental conditions change.

### 2. Active biomass decreases, and lag time increases with depth

As we hypothesized (second hypothesis), relative active biomass declined, and lag time increased with depth (Fig. 5b,c). Reductions in resource availability likely drove these changes in the relative active biomass and lag time (Table 1). Under conditions with relatively low available resources, microbes may enter dormancy (Lennon and Jones, 2011) or inhabit poorly connected colonies without quorum sensing (Mitri et al., 2016), likely increasing microbial lag time. Our results that deep soil layers contain low available resources (Fig. 2) and relatively low active microbes (Fig. 5b), and that lag time quickly drops at a low range of relative active biomass (S3) highlight that any increases in resource availability would disproportionately influence microbial communities in deep soil layers compared to those in topsoil layers. That is, microbial communities in deep soil layers might quickly shorten lag time if increases in resource availability were to drive increases in relative active biomass.

The relative active biomass in this study ranged from 0.02 (=2%) in 0-20 cm soils to 0.0001 (=0.01%) in 210-240 cm soils. Our estimates agree with other studies, where active biomass comprised less than 0.05% (Salazar et al., 2019), 0.1-2% (Blagodatskaya and Kuzyakov, 2013b), 0.2-0.6% (Blagodatskaya et al., 2009), or less than 3.5% (Bloem et al., 1992) of total biomass. These results imply that soil microbial growth is restricted even at topsoil layers with greatest resource availability, possibly due to lack of signal molecules (quorum sensing)(Atkinson and Williams, 2009) or unbalance in stoichiometry between resources and microbial biomass (Keiblinger et al., 2010; Sterner and Elser, 2002). Dormancy is much more common in soil compared to other systems such as freshwater (∼50%) or marine water (∼35%) (Lennon and Jones, 2011). This may be due to heterogeneity in soil’s physicochemical properties and highly fluctuating environmental conditions (Wallenstein and Hall, 2012; Wang et al., 2014). A great proportion of dormancy in soils may help microbial communities cope with patchy and unpredictable environmental variations and sustain their function over longer timescales (Lennon and Jones, 2011).

Many studies demonstrate that active biomass is closely associated with biogeochemical process rates (Salazar-Villegas et al., 2016; Salazar et al., 2019), but we did not detect a clear relationship between active biomass and basal respiration (basal respiration = 0.092 * active biomass + 0.365, R^2^=0.12). One explanation would be extremely low and highly variable basal respiration (0.45 ± 0.02 µg C-CO_2_ g^−1^ dry soil h^−1^; S2) due to relatively low soil OC (Fig. 2a). In other studies, reported basal respiration is higher than ours, at about 180-250 µg C-CO_2_ g^−1^ dry soil h^−1^ at 0-15 cm of soil depth in a temperate forest (Salazar-Villegas et al., 2016) and 3-4 µg C-CO_2_ g^−1^ dry soil h^−1^ from the top 10 cm of a temperate agricultural soil (Blagodatskaya et al., 2014). One study reports comparable values to our estimations at 0.18-0.24 µg C-CO_2_ g^−1^ dry soil h^−1^ in mineral soils (Ritz and Wheatley, 1989). It is plausible that relatively low and variable basal respiration may have decreased our ability to detect differences between vegetation types or among depths, or a relationship between basal respiration and active biomass.

### 3. Deeply rooted plants have little effect on microbial growth parameters

Contrary to our third hypothesis, vegetation types did not influence microbial parameters in this study. The discrepancy between the expectation and the observations is consistent with our finding that switchgrass did not affect C and N availability (Fig. 2). Although the annual cropland was converted to switchgrass more than a decade ago at our study site (Sekaran et al., 2019) and switchgrass developed a deep rooting system (Fig. 1), the stocks of C and N, and ^14^C values of bulk soil under switchgrass were not distinguishable from those under soy (Fig. 2).

Several scenarios might explain the similar resource availability we observed between vegetation types. First, microbial respiration and priming may be greater under switchgrass than soy, canceling out likely higher plant C inputs from more, longer root growth. Soil C stock is a balance between inputs (litter, root exudates, root turnover) and outputs (respiration). If microbial respiration under switchgrass was enhanced (primed) due to greater plant C input, total soil C stocks may be similar between switchgrass and soy. Yet, similar basal respiration between the two vegetation types (S2) and similar ^14^C values in bulk soil OC (Fig. 2b) suggest that this scenario is unlikely. Second, despite more, longer root growth under the switchgrass, the amount of root exudates may be comparable between switchgrass and soy. We did not directly quantify root exudates, but similar DOC concentrations between vegetation types (Fig.2c) suggest that this may be the case. Also, the main function may differ between deep and shallow roots. In a recent review (Lynch, 2019), deep roots are thought to collect water and nitrate, while shallow roots acquire nutrients such as phosphorus, calcium, and potassium. As nitrate is water-soluble, deep roots may not need to release root exudates as shallow roots do to mobilize nutrients. Third, the effects of vegetation and associated changes in the rooting depth on soil C stocks may be minimal in soils with high clay content and alkalinity. Recently, Slessarev et al. (2020b) demonstrated that C stocks were significantly higher under > 10-year-old switchgrass stands relative to annual crops in a low-nutrient sandy soil but found no consistent difference in the amount of C between deeply- and shallow-rooted plants in clayey soil. Soils at our study site were mostly clay loam (approximately 30% clay) with pH higher than 8.0 (S1). Thus, it is plausible that soil conditions may have influenced C accumulation along the soil profiles. Fourth, the annual cropland was fertilized in previous years and soy fixes N from the atmosphere. Contrary to switchgrass, the annual cropland is in rotation between soy and corn, and N fertilizer is used to enhance crop productivity. In addition, soy itself harbors N-fixing bacteria in roots. As such, it is plausible that residual N fertilizer applied in previous years and fixed N may persist to influence resource availability under soy. High dissolved N content under soy along the soil profiles supports this argument (Fig. 2f). Taken together, the conversion from annual crops to switchgrass did not alter resource availability along soil depth profiles at our study site, which, in turn, marginally influenced microbial growth parameters between vegetation types.

## Conclusions

Using a growth kinetics model, we estimated microbial growth parameters throughout soil depth profiles, a key knowledge gap for microbially-driven C dynamics models. While other studies have identified significant differences in profile-scale SOC inventories between switchgrass and shallow rooted conventional crops (Ferchaud et al., 2016; Liebig et al., 2005), in our study site, we did not observe enhanced soil C nor different microbial growth parameters under deep-rooted switchgrass compared to soy. The lack of a detectable plant-cover effect on both SOC stocks and microbial growth capacity may be a function of environmental factors, given that the response of bulk SOC pools to perennial cover can vary considerably as a function of soil properties, environmental conditions, and site history (Blanco-Canqui et al., 2005; Slessarev et al., 2020b). Depth profiles of microbial growth potential revealed increased dormancy rates at depth, but maximum growth potential was relatively similar across soil profiles. This suggests that a component of the microbial community at depth has the potential to rapidly exploit resources introduced by the deep root systems of perennial plants. Thus, to the extent that C inputs from deep root systems can increase SOC stocks, these deep SOC stocks are likely not immune from microbial transformation; rather, they might persist despite the presence of microbial consumers with a high capacity to assimilate fresh C.

## Acknowledgements

We thank Drs. Sandeep Kumar and Udayakumar Sekaram at South Dakota State University for allowing us to collect soil samples in their long-term experimental field site. This work was supported by Lawrence Livermore National Laboratory’s Lab Directed Research and Development program (#19-ERD-010); the National Research Foundation of Korea (NRF) grant funded by the Korea government (MSIT; NRF-2018R1A5A7025409); University of California Merced Chancellor’s Fellowship to KM; University of California Merced and Falasco Endowed Chair to AAB, and work at LLNL was conducted under the auspices of DOE Contract DE-AC52-07NA27344.

## Supplementary figures

**S1.**
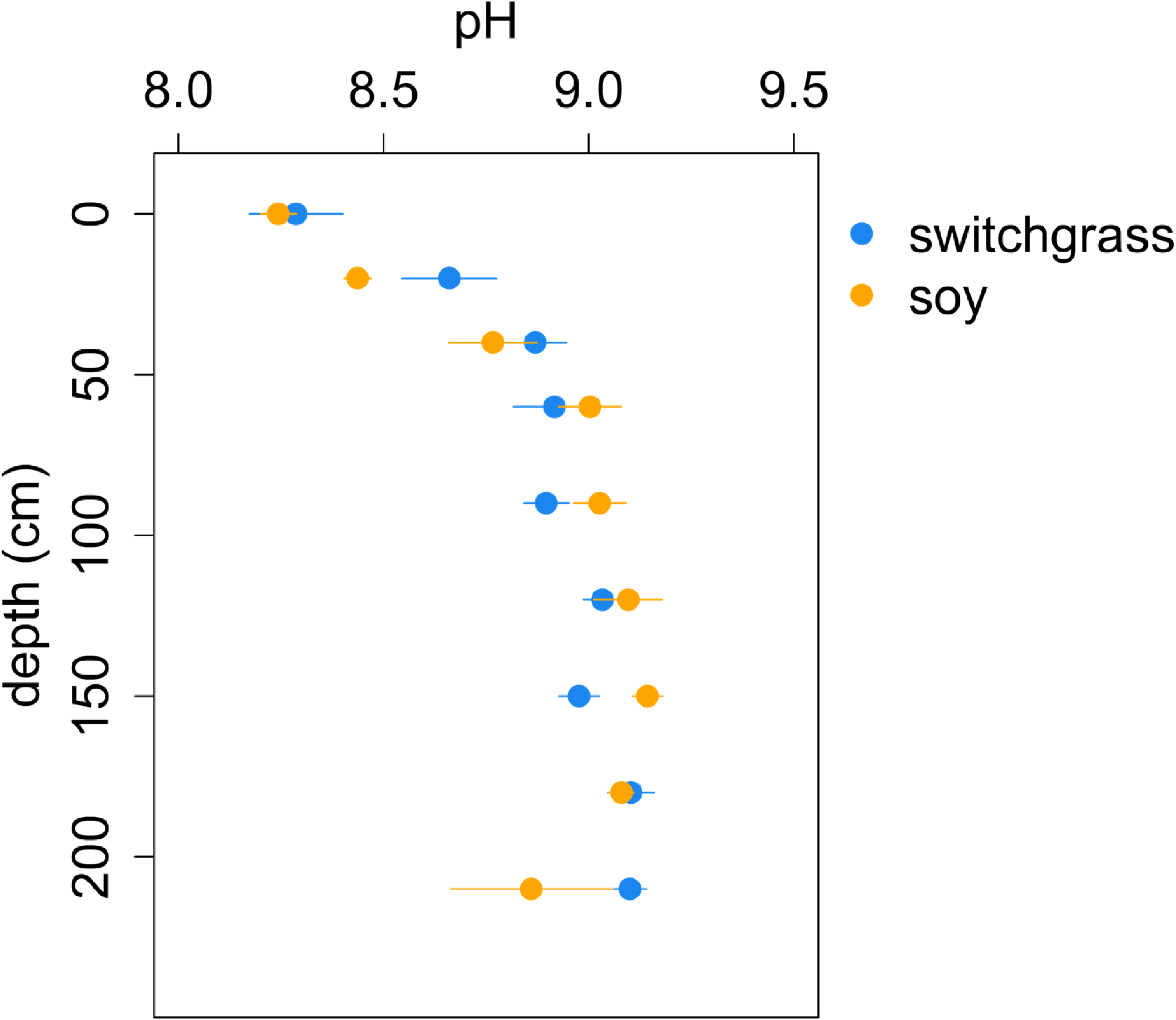
Depth profile of soil pH for switchgrass (blue) and soy (orange) (n=3, error bars represent ± 1 standard error).

**S2.**
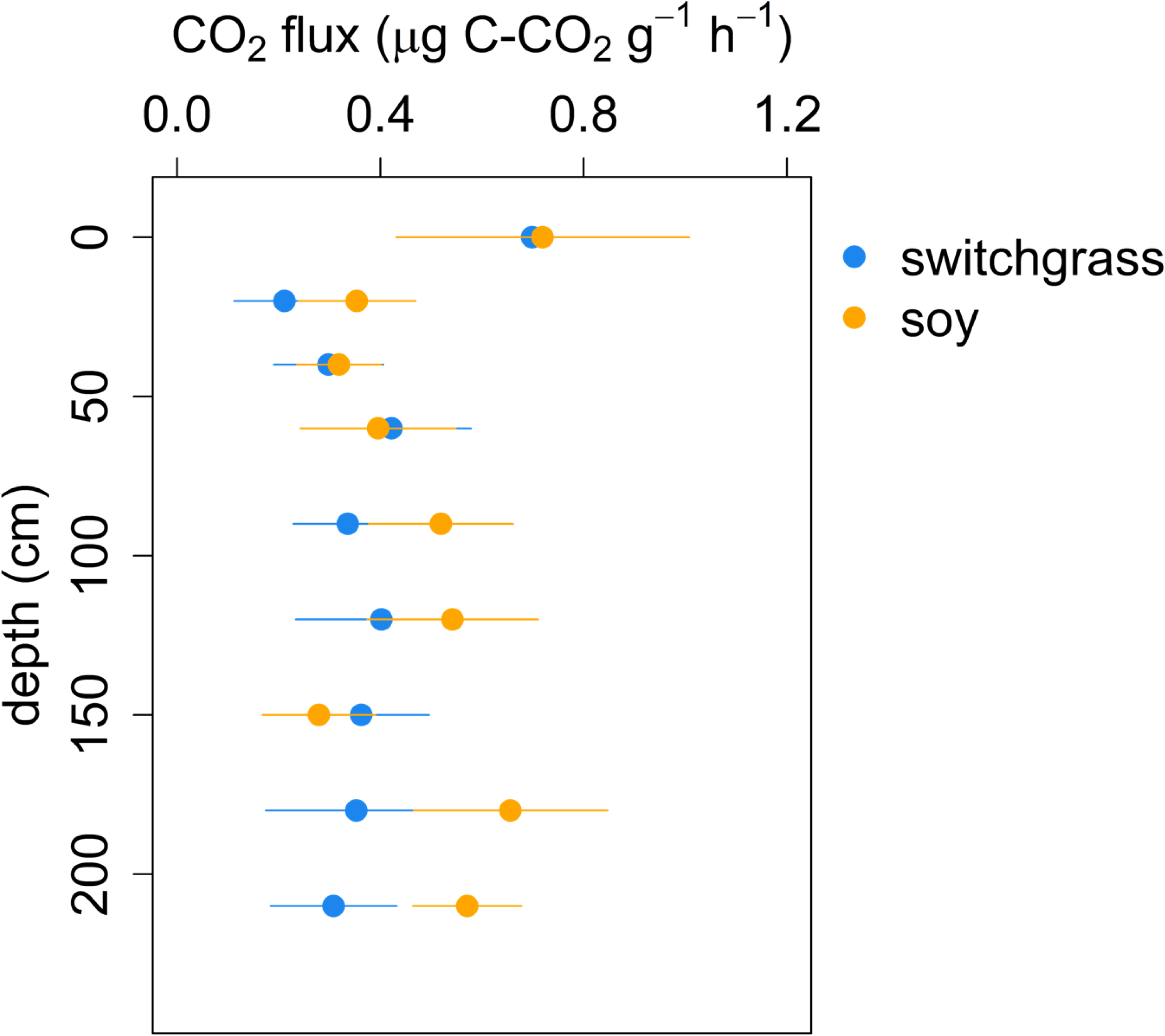
Microbial basal respiration for switchgrass (blue) and soy (orange) soils collected along a 240 cm depth profile from clay-loam agricultural fields in South Dakota (n=3, error bars represent ± 1 standard error).

**S3.**
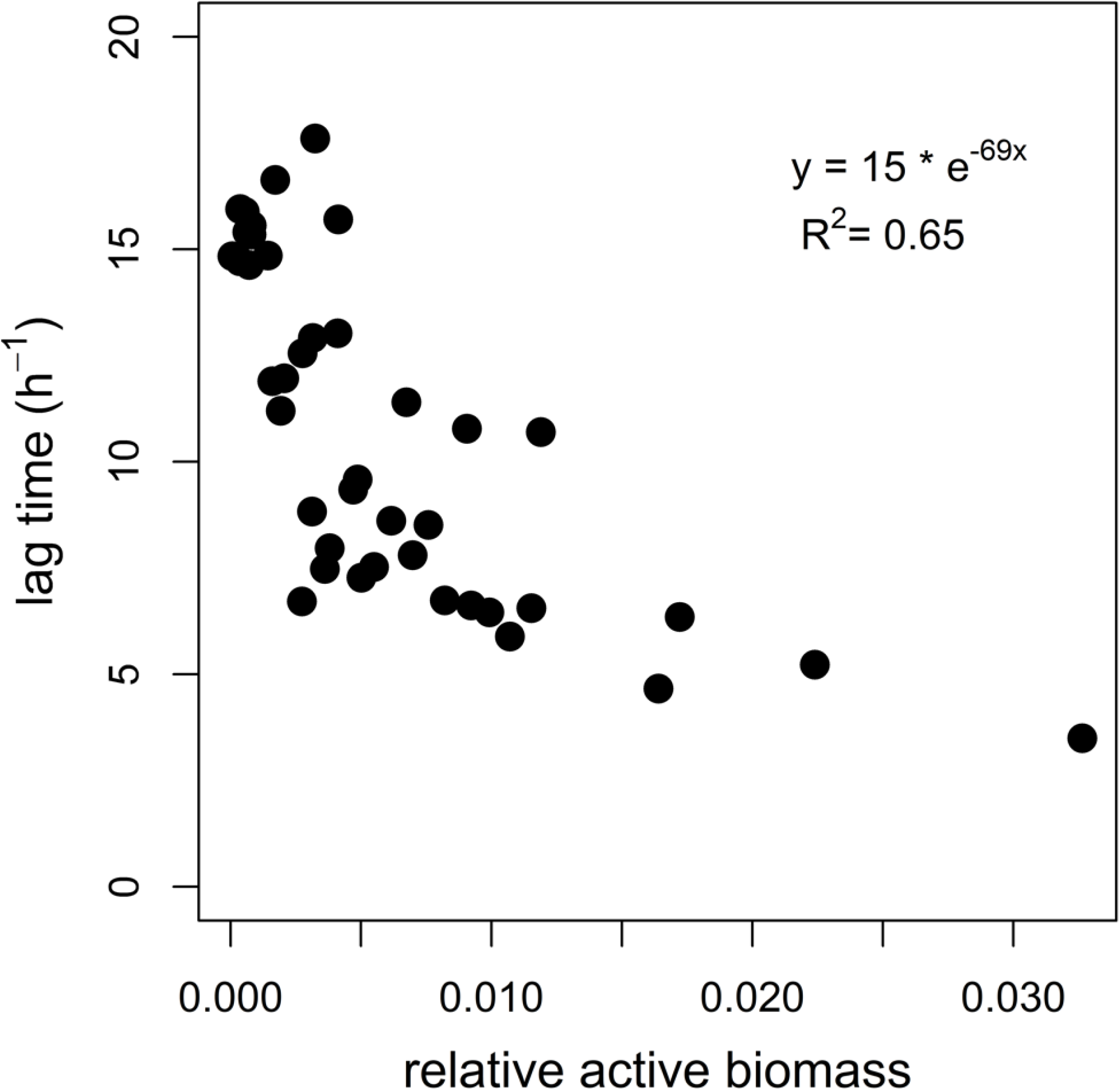
Microbial lag time plotted against relative active biomass across vegetation type and depth.

