## Supplementary figures and images for "Soil depth gradients in microbial growth kinetics under deeply- vs. shallow-rooted plants"

### supplementary

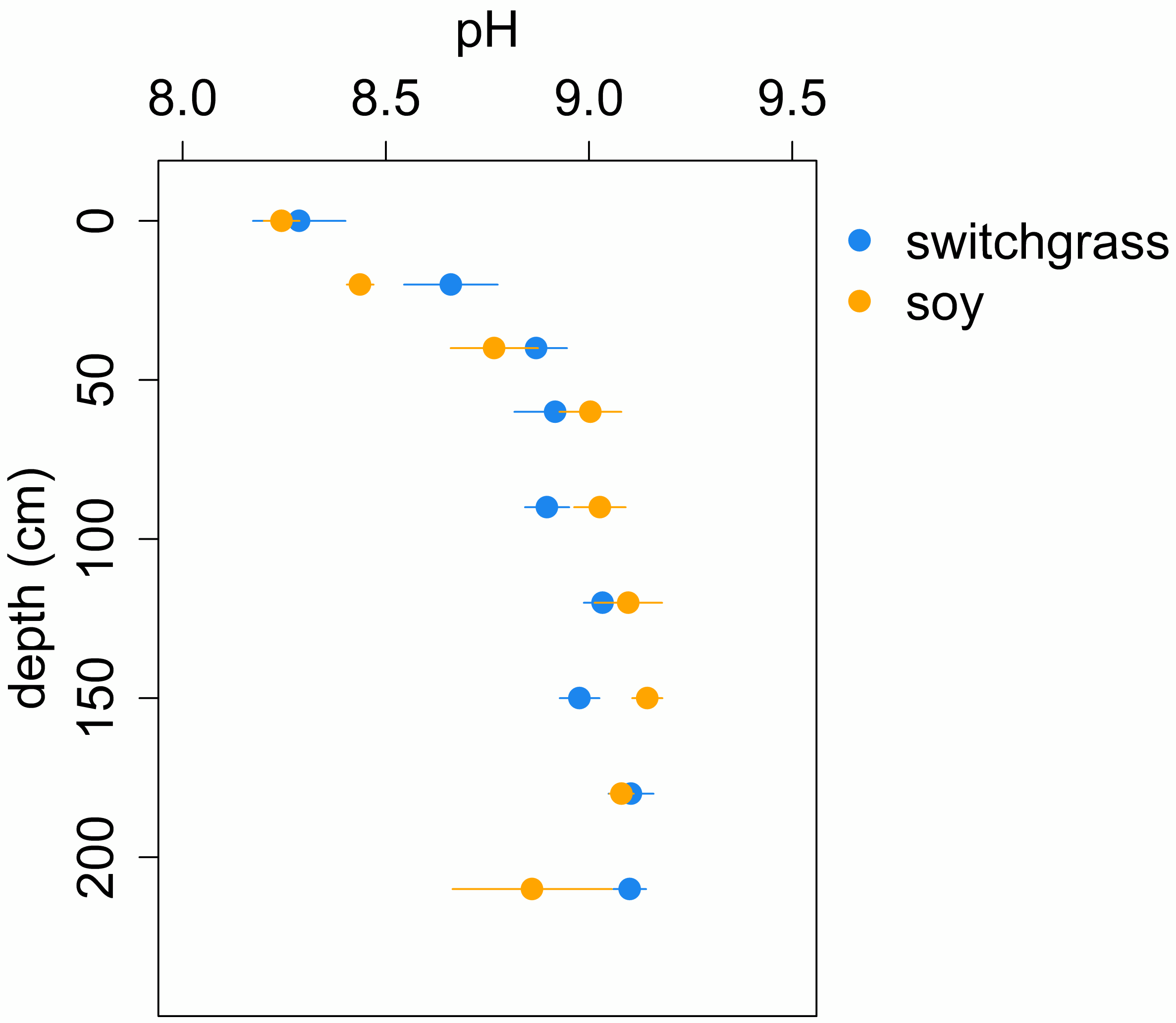

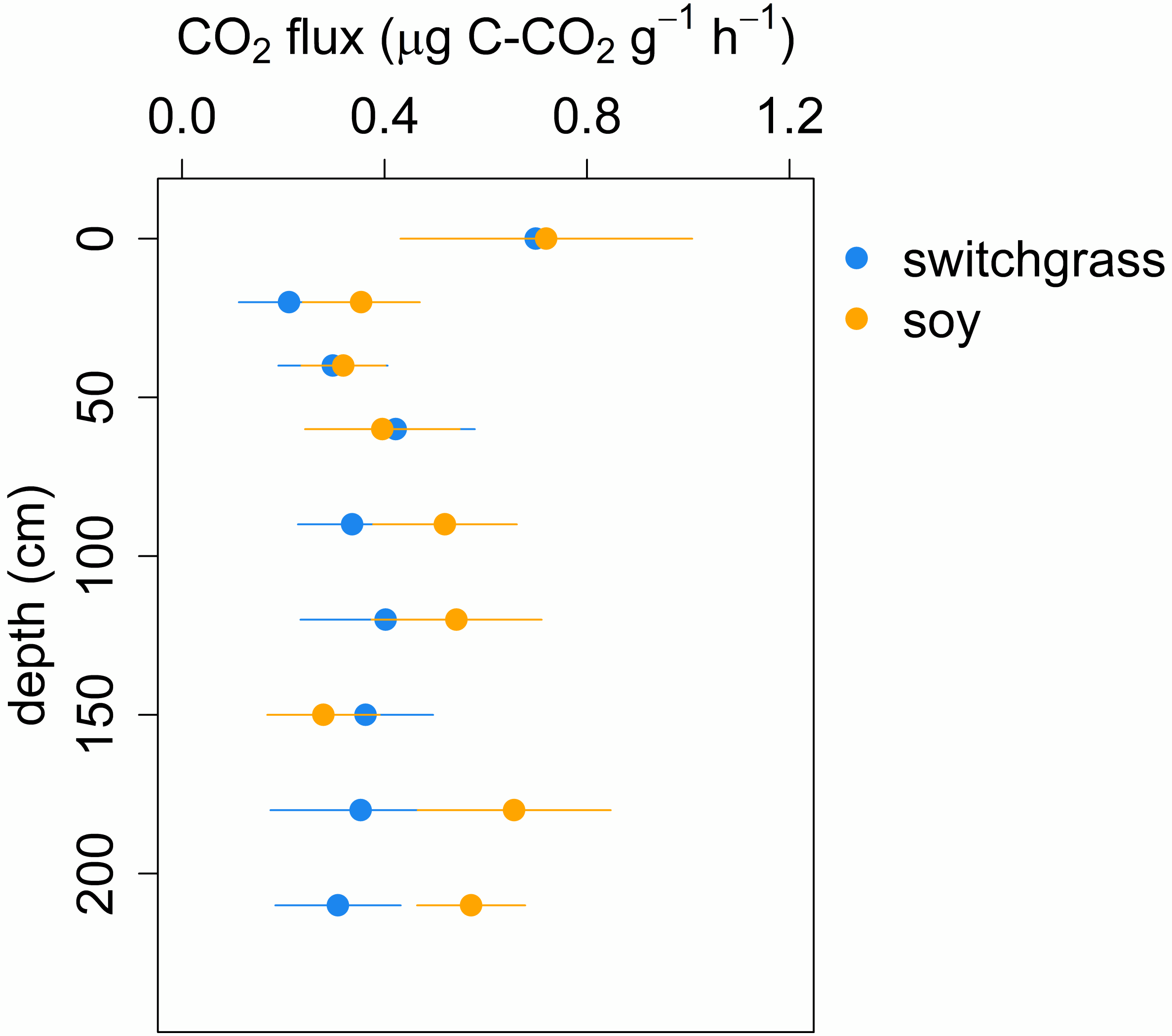

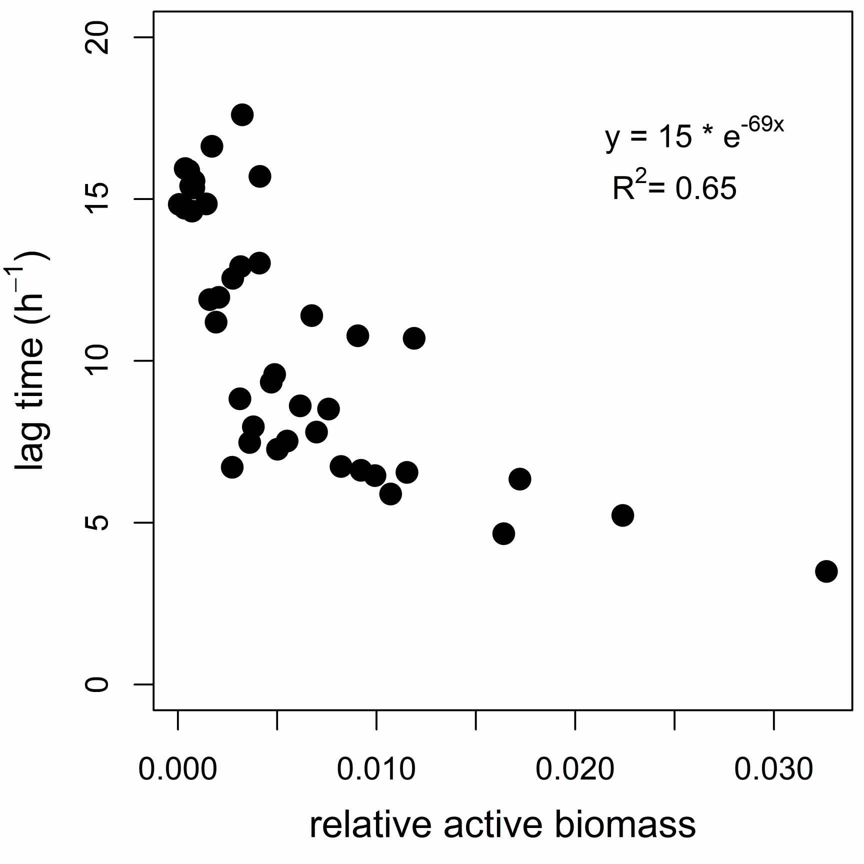
